# Bridging the Gap: Multi-Omics Profiling of Brain Tissue in Alzheimer’s Disease and Older Controls in Multi-Ethnic Populations

**DOI:** 10.1101/2024.04.16.589592

**Authors:** Joseph S. Reddy, Laura Heath, Abby Vander Linden, Mariet Allen, Katia de Paiva Lopes, Fatemeh Seifar, Erming Wang, Yiyi Ma, William L. Poehlman, Zachary S. Quicksall, Alexi Runnels, Yanling Wang, Duc M. Duong, Luming Yin, Kaiming Xu, Erica S. Modeste, Anantharaman Shantaraman, Eric B. Dammer, Lingyan Ping, Stephanie R. Oatman, Jo Scanlan, Charlotte Ho, Minerva M. Carrasquillo, Merve Atik, Geovanna Yepez, Adriana O. Mitchell, Thuy T. Nguyen, Xianfeng Chen, David X. Marquez, Hasini Reddy, Harrison Xiao, Sudha Seshadri, Richard Mayeux, Stefan Prokop, Edward B. Lee, Geidy E. Serrano, Thomas G. Beach, Andrew F. Teich, Varham Haroutunian, Edward J. Fox, Marla Gearing, Aliza Wingo, Thomas Wingo, James J. Lah, Allan I. Levey, Dennis W. Dickson, Lisa L. Barnes, Philip De Jager, Bin Zhang, David Bennett, Nicholas T. Seyfried, Anna K. Greenwood, Nilüfer Ertekin-Taner

## Abstract

**INTRODUCTION:** Multi-omics studies in Alzheimer’s disease (AD) revealed many potential disease pathways and therapeutic targets. Despite their promise of precision medicine, these studies lacked African Americans (AA) and Latin Americans (LA), who are disproportionately affected by AD.

**METHODS:** To bridge this gap, Accelerating Medicines Partnership in AD (AMP-AD) expanded brain multi-omics profiling to multi-ethnic donors.

**RESULTS:** We generated multi-omics data and curated and harmonized phenotypic data from AA (n=306), LA (n=326), or AA *and* LA (n=4) brain donors plus Non-Hispanic White (n=252) and other (n=20) ethnic groups, to establish a foundational dataset enriched for AA and LA participants. This study describes the data available to the research community, including transcriptome from three brain regions, whole genome sequence, and proteome measures.

**DISCUSSION:** Inclusion of traditionally underrepresented groups in multi-omics studies is essential to discover the full spectrum of precision medicine targets that will be pertinent to all populations affected with AD.

## Background

Alzheimer’s disease (AD) is a devastating neurodegenerative disorder that affects millions of people worldwide [1]. While AD is a global health concern, it has been observed that African Americans (AA) and Latin Americans/Latinos/Hispanics (hereafter referred to as LA), are disproportionately affected by the disease [2]. The prevalence of AD in AA is about twice that of non-Hispanic whites (NHW), while LA face a 1.5 times higher risk. Despite these alarming disparities in risk, AA and LA populations remain significantly underrepresented in AD research, including clinical trials, biomarker and genomic studies [3–6].

This underrepresentation becomes even more apparent in genetic studies, where large-scale genome-wide association studies (GWAS) have yielded valuable insights into AD risk factors and potential therapeutic targets. The largest AD GWAS to date comprises over 1 million individuals [7] and, collectively with other studies, identified 75 genetic risk loci [8] in non-Hispanic white (NHW) populations of European ancestry. In contrast, GWAS in AA and LA populations have suffered from limited power [3,4,9–13], with sample sizes less than one to two orders of magnitude of those for NHW populations. Despite limited sample sizes, GWAS and sequencing studies in AA populations identified novel AD risk loci [13–15] and demonstrated allelic heterogeneity for AD risk genes initially discovered in NHW populations, including *TREM2* [14,16] and *ABCA7* [9,14,17]. These findings highlight the potential knowledge to be gained by studying diverse populations to fully capture the genetic and molecular underpinnings of AD in all affected groups. Such knowledge is essential for the development of personalized treatments and interventions for AD using a precision medicine approach like other complex diseases like cancer [18,19].

While genetic variant information is necessary, it is not sufficient to realize the promise of precision medicine. Multi-omics data, including genetic, transcriptome, epigenome, proteome, metabolome, and lipidome data, generated and analyzed in large-scale, diverse, and deeply phenotyped individuals, are required to uncover disease pathways and mechanisms in all affected populations. Thus, novel personalized therapies and biomarkers can be attainable by deciphering the complex molecular etiopathogenesis of AD. With a goal to accelerate discovery of candidate drug targets and translate these discoveries to new therapies for AD, the Accelerating Medicines Partnership in AD (AMP-AD) Target Discovery and Preclinical Validation Project was launched in 2014 bringing together NIA-supported foundational grants [20]. This effort led to the generation and analysis of RNA-sequencing (RNAseq) based transcriptome, whole genome sequence (WGS), proteome, metabolome, and epigenome data on more than 2,500 brain samples primarily from NHW donors with AD and non-AD neuropathologies, as well as unaffected controls. This vast amount of data has been made available to the research community [21–24] simultaneously with data quality control (QC) and without publication embargoes and can be accessed through the AD Knowledge Portal [20,25]. These rich, high-quality data have been utilized to identify or validate potential risk mechanisms in AD and other neurodegenerative diseases (examples include [22–24,26–53]) and led to the data-driven identification and nomination of over 600 key driver genes/candidate targets for AD. These target nominations and the associated data, including a set of curated genomic analyses and information on their druggability, have been made available via the AMP-AD open-source platform Agora (https://agora.ampadportal.org/).

Despite these advances, such multi-omics studies of AD and related disorders (ADRD) have lacked sampling from AA and LA populations with few exceptions [54,55]. To bridge the data and knowledge gap in multi-omics studies of underrepresented populations in AD research, AMP-AD investigators launched a diversity initiative to expand molecular profiling of brain tissue to multi-ethnic donors. We generated whole genome sequence (WGS), transcriptome, and proteome data; curated and harmonized phenotypic data from AA (n=306), LA (n=326), and AA and LA (4) brain donors as well as NHWs (n=252) and other (n=20) ethnic groups to establish a foundational multi-omics dataset enriched for AA and LA participants. This study describes this unique dataset made available to the research community. These data will lay the groundwork for bridging the knowledge disparities in AD research and are expected to uncover pathways, molecules, and genetic variants that drive or contribute to AD in these populations. By focusing on these high-risk populations and leveraging the infrastructure developed by AMP-AD, this initiative promotes inclusivity in research, is aligned with the broader goal of advancing precision medicine for *All of AD* in the spirit of the National Institutes of Health *All of Us* program [56] and aims to ultimately improve lives of all individuals affected by this devastating disease.

## Methods and Results

### Study populations by biospecimen and data-contributing institutions

Five AMP-AD data contributing institutions participated in providing brain samples and associated data for the AMP-AD Diversity Initiative, which is enriched for donors from AA and LA populations. Each of the following institutions (Mayo Clinic, Rush University, Mount Sinai, Columbia University, and Emory University) coordinated the collection of these brain samples from their own networks of affiliated brain banks, cohort studies, and Alzheimer’s Disease Research Centers (**Table 1**). In addition to new donors from the studies and cohorts described below, 303 predominantly NHW (96%) individuals previously characterized in the AMP-AD 1.0 initiative are described in **Supplementary Table 1**. Two-hundred eighty-four samples from these individuals were included in the proteomics to provide more balance to the samples (described in Methods). The other 19 samples have newly generated transcriptomic or proteomic data as part of the Diverse Cohorts initiative but only have WGS available from AMP-AD 1.0.

**Table 1.**
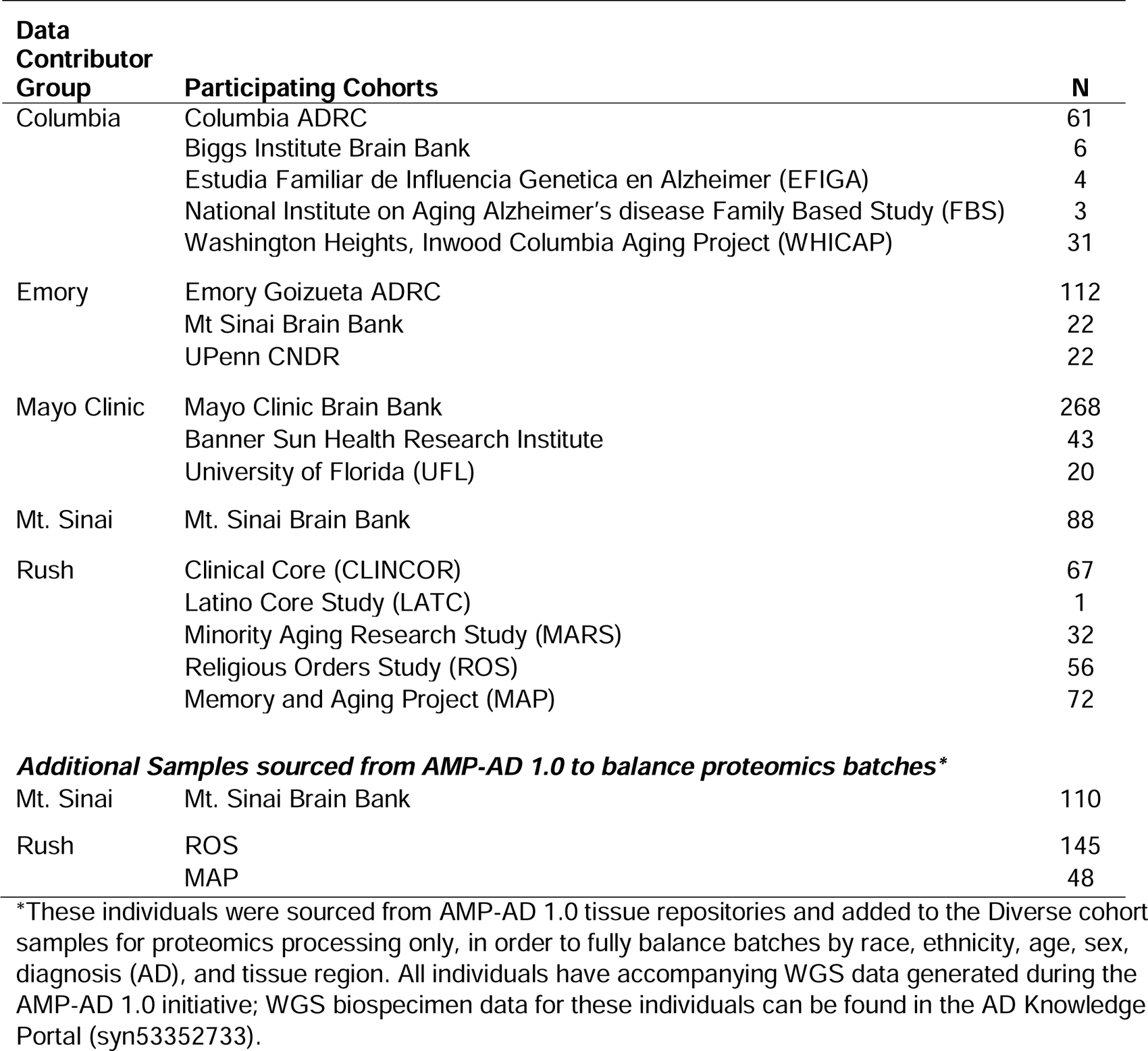
Tissue sample sources by contributing institutions and cohorts.

| **Data Contributor Group** | **Participating Cohorts** | **N** |
| --- | --- | --- |
| Columbia | Columbia ADRC | 61 |
|  | Biggs Institute Brain Bank | 6 |
|  | Estudia Familiar de Influenza Genetica en Alzheimer (EFIGA) | 4 |
|  | National Institute on Aging Alzheimer's disease Family Based Study (FBS) | 3 |
|  | Washington Heights, Inwood Columbia Aging Project (WHICAP) | 31 |
| Emory | Emory Goizueta ADRC | 112 |
|  | Mt Sinai Brain Bank | 22 |
|  | UPenn CNDR | 22 |
| Mayo Clinic | Mayo Clinic Brain Bank | 268 |
|  | Banner Sun Health Research Institute | 43 |
|  | University of Florida (UFL) | 20 |
| Mt. Sinai | Mt. Sinai Brain Bank | 88 |
| Rush | Clinical Core (CLINCOR) | 67 |
|  | Latino Core Study (LATC) | 1 |
|  | Minority Aging Research Study (MARS) | 32 |
|  | Religious Orders Study (ROS) | 56 |
|  | Memory and Aging Project (MAP) | 72 |
| ***Additional Samples sourced from AMP-AD 1.0 to balance proteomics batches\**** |  |  |
| Mt. Sinai | Mt. Sinai Brain Bank | 110 |
| Rush | ROS | 145 |
|  | MAP | 48 |

\*These individuals were sourced from AMP-AD 1.0 tissue repositories and added to the Diverse cohort samples for proteomics processing only, in order to fully balance batches by race, ethnicity, age, sex, diagnosis (AD), and tissue region. All individuals have accompanying WGS data generated during the AMP-AD 1.0 initiative; WGS biospecimen data for these individuals can be found in the AD Knowledge Portal (syn53352733).

#### Mayo Clinic

Brain samples provided by the Mayo Clinic were from three brain banks: Mayo Clinic Florida Brain Bank (n=268), the Arizona Study of Aging and Neurodegenerative Disorders and Brain and Body Donation Program at Banner Sun Health (n=43), and the University of Florida Human Brain and Tissue Bank (n=20). There were 53 AA, 182 LA, and 96 NHW brain donors. Tissue samples from the superior temporal gyrus, anterior caudate nucleus, and dorsolateral prefrontal cortex were obtained from the donors. The Mayo Clinic Institutional Review Board approved all of this work. All donors or their next of kin provided informed consent.

***Mayo Clinic Brain Bank*** collects brain specimens with neurodegenerative diseases as well as unaffected controls. All donors from the Mayo Clinic Brain Bank underwent neuropathologic evaluation by Dr. Dennis W. Dickson. Neuropathologic AD diagnosis was made according to the NINCDS-ADRDA criteria [57], such that all AD donors had Braak neurofibrillary tangle (NFT) stage of IV or greater and evidence of Thal 2 or greater amyloid deposits.

**The Arizona Study of Aging and Neurodegenerative Disorders and Brain and Body Donation Program at Banner Sun Health (Banner)** has collected brains and whole body donations since 1987 [58]. Donors are residents of retirement communities in Phoenix, Arizona, and are typically enrolled when they are cognitively normal, with directed recruitment efforts are aimed at individuals with AD, Parkinson’s Disease, and cancer. Neuropathological diagnosis of AD followed standard NIA guidelines [59].

***University of Florida (UFL)*** samples were collected through the University of Florida Human Brain and Tissue Bank (UF HBTB). All University of Florida brains underwent neuropathological diagnosis of AD according to current NIA guidelines [59,60], with any degree of AD neuropathologic change resulting in an AD diagnosis.

#### Emory University

All samples were collected as part of ongoing studies at Emory’s Goizueta Alzheimer’s Disease Research Center (ADRC), including participants in the ADRC Clinical Core, the Emory Healthy Brain Study, and the ADRC-affiliated Emory Cognitive Neurology Clinic. AD cases were consistent with NIA-Reagan criteria for “High Likelihood” [61]. In addition, investigators at Emory reviewed banked tissue samples previously sent to Emory as part of the AMP-AD 1.0 initiative (but never were submitted for -omics generation until now) and included tissues from the University of Pennsylvania Integrated Neurodegenerative Disease Brain Bank [62] and Mount Sinai Brain Bank [23] to maximize the number of AA and LA samples and provide balance in their proteomics batching (as described in Methods). There were 75 AA, 5 LA, and 76 NHW donors with new data generated as part of the Diverse Cohorts initiative. Further, 284 samples with transcriptomics and/or WGS data generated as part of the AMP-AD 1.0 initiative were added to provide further balance to proteomics batching. Tissue samples were obtained from the anterior caudate nucleus, dorsolateral prefrontal cortex, and the superior temporal gyrus. All participants provided informed consent under protocols approved by Emory University’s Institutional Review Board.

#### Rush University

Multiple longitudinal, epidemiologic cohort studies of aging and the risk of AD are conducted by Rush Alzheimer’s Disease Center (RADC) and include Clinical Core (CLINCOR), Latino Core Study (LATC), Minority Aging Research Study (MARS), Religious Orders Study (ROS), and Memory Aging Project (MAP). Most of the participants of these cohorts are older adults aged 65 and above, encompassing a range of ethnic and demographic backgrounds. They do not have known dementia at enrollment and agree to undergo annual clinical evaluations, with optional brain donation. There were 113 AA, 45 LA, 11 Asian, 49 NHW, 1 American Indian or Alaska Native, 4 American Indian or Alaska Native donors who also identified as Hispanic, and 3 AA donors who also identified as Hispanic. Tissue samples were obtained from the anterior caudate nucleus, dorsolateral prefrontal cortex, and the superior temporal gyrus. Informed consent and IRB approvals were obtained under the Rush University IRB. Details for each cohort are as follows:

***Clinical Core (CLINCOR)*** studies the transition from normal aging to mild cognitive impairment (MCI) to the earliest stages of dementia. Enrollment started in 1992, primarily with individuals diagnosed with dementia. Since 2008, the study has transitioned to consist of primarily AA, most without dementia, who share a common core of risk factors with the other RADC studies. The participants are from the metropolitan Chicago area and outlying suburbs.

***Latino Core Study (LATC)*** is a cohort study of cognitive decline aiming to identify risk factors of AD in older Latinos. The participants self-identified as Latino/Hispanic, and enrollment started in 2015. Recruitment locations include churches, subsidized senior housing facilities, retirement communities, Latino/Hispanic clubs, organizations, and social service centers that cater to seniors in various Chicago neighborhoods and outlying suburbs.

***Minority Aging Research Study (MARS)*** is a cohort study of cognitive decline and risk of AD in older AAs. The recruitment began in 2004, and brain donation in 2010. The participants were recruited from various places, including churches, senior housing facilities, retirement communities, AA clubs, organizations, fraternities and sororities, and social service centers catering to seniors in metropolitan Chicago and outlying suburbs [63].

***Religious Orders Study (ROS)*** *and **Memory and Aging Project (MAP)*** are prospective community-based studies of risk factors for cognitive decline, incident AD dementia, and other health outcomes. ROS began to recruit catholic nuns, priests, and brothers from across the United States in 1994. MAP started recruiting participants from retirement communities and subsidized senior housing facilities throughout Chicago and northeastern Illinois in 1997 [64]. The ROSMAP participants are primarily non-Latino White, with small proportions of AA, Latino, and other racial groups.

#### Mount Sinai School of Medicine (MSSM)

The MSSM cohort comprises donor brain tissue obtained from the Mount Sinai/JJ Peters VA Medical Center Brain Bank (MSBB) [23,65]. There were 31 AA, 27 LA, and 30 NHW donors. Tissue samples were obtained from the anterior caudate nucleus, dorsolateral prefrontal cortex, and superior temporal gyrus. Autopsy protocols were approved by the Mount Sinai and JJ Peters VA Medical Center Institutional Review Boards, and patient consent for donation was obtained.

#### Columbia University

Samples were collected from the New York Brain Bank (NYBB) at Columbia University, which was established to collect postmortem human brains to further study neurodegenerative disorders. There were 35 AA donors (one also identified as LA), 68 LA, 1 NHW, and 1 Asian donor. Tissue samples were obtained from the anterior caudate nucleus, dorsolateral/dorsomedial prefrontal cortex, and temporal pole. The appropriate review boards approved this study. The brain tissues contributed by Columbia University come from the following cohorts, brain banks, and studies:

***The Columbia Alzheimer’s Disease Research Center (Columbia ADRC)*** cohort consists of clinical participants in the Columbia ADRC who agreed to brain donation. All banked brains have one hemisphere fixed for subsequent diagnostic evaluation, and one hemisphere is banked fresh. For the hemisphere that is banked fresh, we block and freeze regions that are most commonly requested by researchers using liquid nitrogen, and specimens are stored at -80 °C. This is performed on all ADRC brain donations, as well as on brains from the additional cohorts described below that also contributed to this study.

***National Institute on Aging Alzheimer’s Disease Family Based Study (NIA-AD FBS)*** has recruited and followed 1,756 families with suspected late-onset Alzheimer’s Disease (AD), including 9,682 family members and 1,096 unrelated, nondemented elderly from different racial and ethnic groups. This Resource Related Cooperative Agreement has now extended to the recruitment of familial early-onset AD. The goals of this protocol are to provide rich genetic and biological resources for the scientific community, which includes longitudinal phenotype data, genotyped data, as well as brain tissue, plasma, and PBMCs.

***Washington Heights, Inwood Columbia Aging Project (WHICAP)*** includes representative proportions of AA (28%), Caribbean Hispanics (48%), and non-Hispanic whites (24%). Since its inception in 1992, over 6,000 participants have enrolled in this Program Project. Over the length of the project, we have identified environmental, health-related, and genetic risk factors of disease and predictors of disease progression by collecting longitudinal data on cognitive performance, emotional health, independence in daily activities, blood pressure, anthropometric measures, cardiovascular status and selected biomarkers in this elderly, multi-ethnic cohort. WHICAP includes Biomarker studies, MRI, PET scans, and brain tissue.

***The Biggs Institute Brain Bank*** at the University of Texas Health Science Center at San Antonio is a biorepository and research laboratory focused on the pathology of neurodegenerative disorders in the San Antonio metropolitan region and the greater South Texas. The Biggs Institute Brain Bank is the central service provider for the South Texas Alzheimer’s Disease Research Center Neuropathology Core, collecting postmortem brain, spinal cord, cerebrospinal fluid, and dermal tissue from study participants and donors. Brain donation consent was obtained from the donor’s legal next-of-kin prior to the autopsy. Autopsied brain tissue is hemisected, with the left hemibrain (typically) fixed in 10% neutral-buffered formalin and the right hemibrain (typically) sectioned fresh and preserved at -80°C. Following a minimum 4-week fixation period and postmortem *ex vivo* magnetic resonance imaging [66], fixed tissue is sectioned and sampled in accordance with National Institute on Aging-Alzheimer’s Association Alzheimer’s disease (AD) neuropathologic guidelines. For the AMP-AD Diversity Initiative, frozen tissue (approximately 500 mg) was sampled from the anterior caudate, the middle frontal gyrus (Brodmann Area 9 or dorsolateral prefrontal cortex; at the same level as the anterior caudate), and the superior temporal gyrus (at the level of the amygdala) from 6 brain autopsy cases in the Biggs Institute Brain Bank. All research and tissue-sharing activities herein were reviewed and approved by the University of Texas Health Science Center at San Antonio Institutional Review Board and Office of Sponsored Projects.

***Estudio Familiar de Influencia Genetica en Alzheimer (EFIGA)*** is a family-based study initiated in 1998. The study included 683 at-risk family members from 242 AD-affected families of Caribbean Hispanic descent, recruited from clinics in the Dominican Republic and the Taub Institute on Alzheimer’s Disease and the Aging Brain in New York. An AD case was defined as any individual meeting NINCDS-ADRD criteria [57] for probable or possible late-onset Alzheimer’s Disease (LOAD).

### Demographic, clinical, and neuropathologic variables collected

Each donor with brain samples included in the AMP-AD Diversity Initiative was assigned a non-identifiable individual ID by the contributing institution. For each participant, the same demographic variables were curated: cohort (or initial study group population to which the participant belonged); sex (male or female); self-reported race (American Indian or Alaska Native, Asian, Black or African American, White, Other); self-reported ethnicity (a true/false indicator for “is Latin American/Hispanic”); age of death in years (individuals 90 and over were designated as “90+” according to HIPAA privacy rules); post-mortem interval in hours where available; and *APOE* genotype.

The results of standard neuropathological assessments previously performed on the donor brains were also collected from the relevant brain banks and harmonized when possible, following the harmonization protocols established by the Alzheimer’s Disease Sequencing Project Phenotype Harmonization Consortium, as noted in their Neuropathology data dictionary (https://vmacdata.org/adsp-phc). Post-mortem Thal amyloid stages [67] were available for Mayo Clinic, Emory, and a subset of Rush donors. All other donors were assigned a semi-quantitative measure of neuritic plaque on a four-point scale, the Consortium to Establish a Registry for Alzheimer’s Disease (CERAD) score [68]. A semiquantitative measure of the severity of neurofibrillary tangle pathology, Braak Stage (values equal to 0, I, II, III, IV, V, or VI) was included for all donors [69].

### Donor characteristics

Donor characteristics varied by the contributing institution (**Table 2**). The overall median age of all participants was 82 years old, with 88.9% of the participants age 65 and older. A larger proportion of participants were female (59.0%) than male.

**Table 2.**
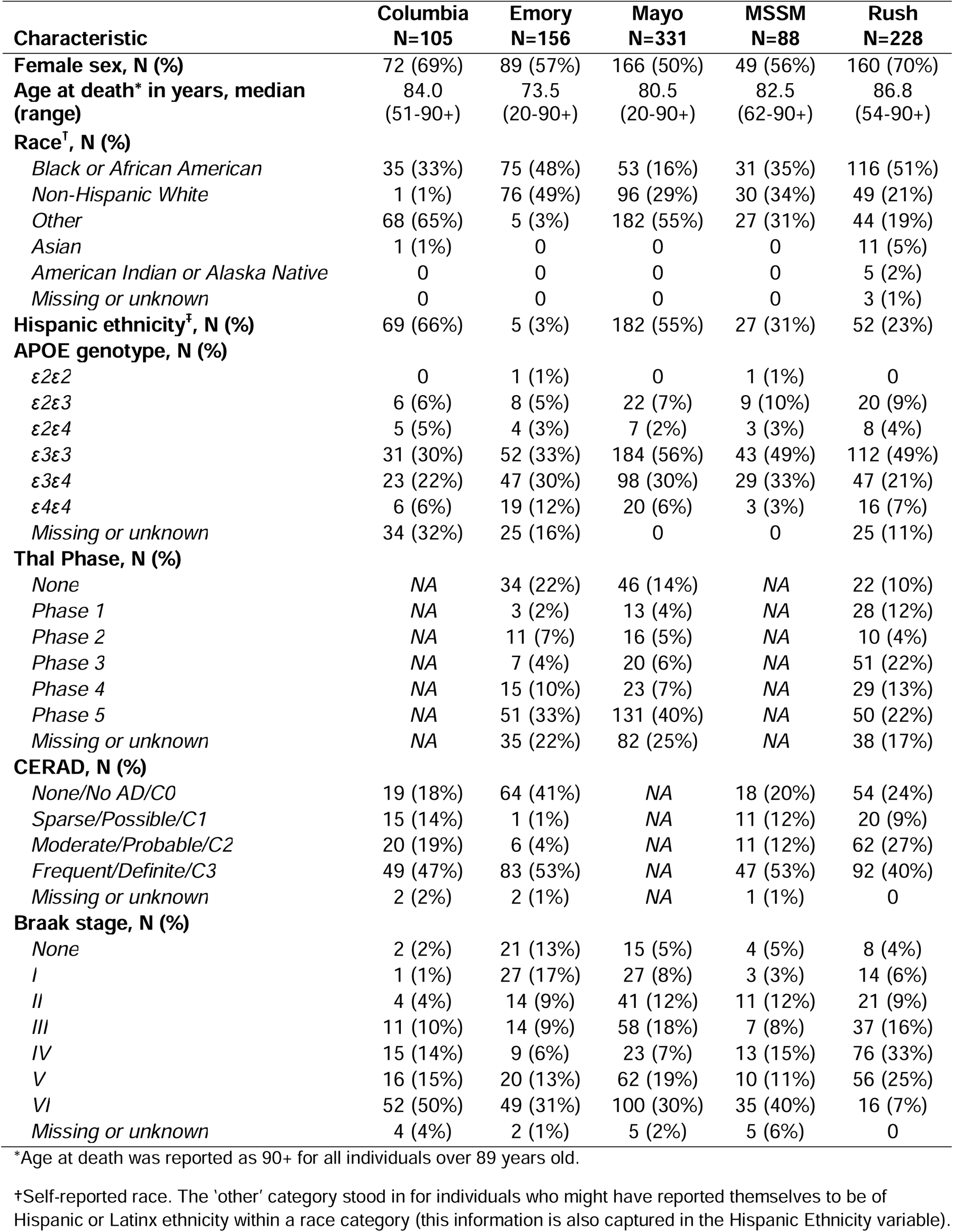
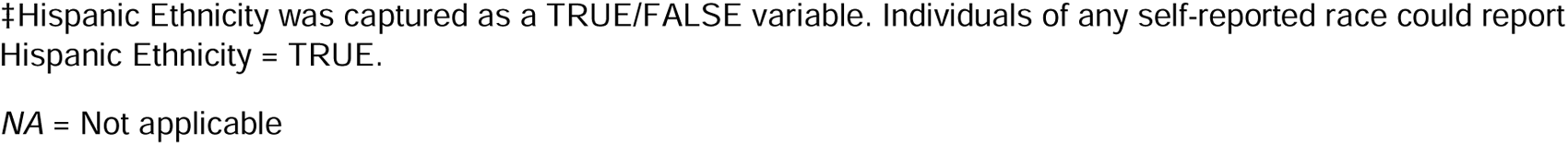
Donor Characteristics by Contributing Institution.

| Characteristic | Columbia<br>N=105 | Emory<br>N=156 | Mayo<br>N=331 | MSSM<br>N=88 | Rush<br>N=228 |
| --- | --- | --- | --- | --- | --- |
| **Female sex, N (%)** | 72 (69%) | 89 (57%) | 166 (50%) | 49 (56%) | 160 (70%) |
| **Age at death\* in years, median (range)** | 84.0<br>(51-90+) | 73.5<br>(20-90+) | 80.5<br>(20-90+) | 82.5<br>(62-90+) | 86.8<br>(54-90+) |
| **Race<sup>†</sup>, N (%)** |  |  |  |  |  |
| Black or African American | 35 (33%) | 75 (48%) | 53 (16%) | 31 (35%) | 116 (51%) |
| Non-Hispanic White | 1 (1%) | 76 (49%) | 96 (29%) | 30 (34%) | 49 (21%) |
| Other | 68 (65%) | 5 (3%) | 182 (55%) | 27 (31%) | 44 (19%) |
| Asian | 1 (1%) | 0 | 0 | 0 | 11 (5%) |
| American Indian or Alaska Native | 0 | 0 | 0 | 0 | 5 (2%) |
| Missing or unknown | 0 | 0 | 0 | 0 | 3 (1%) |
| **Hispanic ethnicity<sup>†</sup>, N (%)** | 69 (66%) | 5 (3%) | 182 (55%) | 27 (31%) | 52 (23%) |
| **APOE genotype, N (%)** |  |  |  |  |  |
| ε2ε2 | 0 | 1 (1%) | 0 | 1 (1%) | 0 |
| ε2ε3 | 6 (6%) | 8 (5%) | 22 (7%) | 9 (10%) | 20 (9%) |
| ε2ε4 | 5 (5%) | 4 (3%) | 7 (2%) | 3 (3%) | 8 (4%) |
| ε3ε3 | 31 (30%) | 52 (33%) | 184 (56%) | 43 (49%) | 112 (49%) |
| ε3ε4 | 23 (22%) | 47 (30%) | 98 (30%) | 29 (33%) | 47 (21%) |
| ε4ε4 | 6 (6%) | 19 (12%) | 20 (6%) | 3 (3%) | 16 (7%) |
| Missing or unknown | 34 (32%) | 25 (16%) | 0 | 0 | 25 (11%) |
| **Thal Phase, N (%)** |  |  |  |  |  |
| None | NA | 34 (22%) | 46 (14%) | NA | 22 (10%) |
| Phase 1 | NA | 3 (2%) | 13 (4%) | NA | 28 (12%) |
| Phase 2 | NA | 11 (7%) | 16 (5%) | NA | 10 (4%) |
| Phase 3 | NA | 7 (4%) | 20 (6%) | NA | 51 (22%) |
| Phase 4 | NA | 15 (10%) | 23 (7%) | NA | 29 (13%) |
| Phase 5 | NA | 51 (33%) | 131 (40%) | NA | 50 (22%) |
| Missing or unknown | NA | 35 (22%) | 82 (25%) | NA | 38 (17%) |
| **CERAD, N (%)** |  |  |  |  |  |
| None/No AD/C0 | 19 (18%) | 64 (41%) | NA | 18 (20%) | 54 (24%) |
| Sparse/Possible/C1 | 15 (14%) | 1 (1%) | NA | 11 (12%) | 20 (9%) |
| Moderate/Probable/C2 | 20 (19%) | 6 (4%) | NA | 11 (12%) | 62 (27%) |
| Frequent/Definite/C3 | 49 (47%) | 83 (53%) | NA | 47 (53%) | 92 (40%) |
| Missing or unknown | 2 (2%) | 2 (1%) | NA | 1 (1%) | 0 |
| **Braak stage, N (%)** |  |  |  |  |  |
| None | 2 (2%) | 21 (13%) | 15 (5%) | 4 (5%) | 8 (4%) |
| I | 1 (1%) | 27 (17%) | 27 (8%) | 3 (3%) | 14 (6%) |
| II | 4 (4%) | 14 (9%) | 41 (12%) | 11 (12%) | 21 (9%) |
| III | 11 (10%) | 14 (9%) | 58 (18%) | 7 (8%) | 37 (16%) |
| IV | 15 (14%) | 9 (6%) | 23 (7%) | 13 (15%) | 76 (33%) |
| V | 16 (15%) | 20 (13%) | 62 (19%) | 10 (11%) | 56 (25%) |
| VI | 52 (50%) | 49 (31%) | 100 (30%) | 35 (40%) | 16 (7%) |
| Missing or unknown | 4 (4%) | 2 (1%) | 5 (2%) | 5 (6%) | 0 |

\*Age at death was reported as 90+ for all individuals over 89 years old.
†Self-reported race. The 'other' category stood in for individuals who might have reported themselves to be of Hispanic or Latinx ethnicity within a race category (this information is also captured in the Hispanic Ethnicity variable).
‡Hispanic Ethnicity was captured as a TRUE/FALSE variable. Individuals of any self-reported race could report Hispanic Ethnicity = TRUE. NA = Not applicable

### Diagnostic harmonization

AMP-AD Diversity Initiative has brain biospecimens from archival brain banks (e.g., Mayo Clinic) and from participants who were followed clinically while living before they came to autopsy (e.g., Rush University ROS and MAP cohorts). Donors from archival brain banks may not have a clinical diagnosis, while all donors had neuropathologic variables that enabled neuropathologic diagnosis. Since cohorts had variable clinical and neuropathological diagnostic information regarding AD case status, we chose to determine AD case/control status according to neuropathologic data for purposes of cross-cohort analysis (**Table 3**). For all individuals with measures of CERAD and Braak, we calculated a modified NIA Reagan diagnosis of AD [61], resulting in the following outcomes: No AD, Low Likelihood of AD, Intermediate Likelihood of AD, and High likelihood of AD. Mayo Clinic Brain Bank donors, which constituted the largest overall and single brain bank group contributing to the AMP-AD Diversity Initiative, lacked CERAD scores but had AD diagnoses according to NINCDS-ADRDA criteria [57]. Mayo Clinic Brain Bank donors were diagnosed as definite AD if they had Braak Stage greater than or equal to IV and the presence of amyloid beta plaques as assessed by a single neuropathologist (Dr. Dennis W. Dickson). Mayo Clinic Brain Bank donors were diagnosed as controls if they had Braak Stage less than or equal to III, sparse or no Aß plaques, and lacked any other neuropathologic diagnosis for neurodegenerative diseases. For all donors, we established the following criteria to achieve a uniform neuropathologic diagnosis of AD and to harmonize AD case/control diagnoses between cohorts as closely as possible: AD diagnosis was assigned to individuals with Braak Stage ≥ IV and CERAD measure equal to Moderate/Probable AD or Frequent/Definite AD. Control diagnosis was assigned to individuals with Braak stage ≤ III and CERAD measure equal to None/No AD or Sparse/Possible AD. Any donors who did not fall under these criteria were assigned as ‘Other.’ These thresholds, while imperfect, are relatively conservative and also serve to exclude individuals with age-related tauopathies from having an AD case or control designation.

**Table 3.** Neuropathologic Diagnoses by Contributing Institution.

| Outcome | Columbia<br>N=105 | Emory<br>N=156 | Mayo<br>N=331 | MSSM<br>N=88 | Rush<br>N=228 |
| --- | --- | --- | --- | --- | --- |
| **NIA Reagan\*** |  |  |  |  |  |
| No AD | 2 (2%) | 21 (13%) | NA | 4 (5%) | 4 (2%) |
| Low Likelihood | 30 (29%) | 45 (29%) | NA | 24 (27%) | 78 (34%) |
| Intermediate Likelihood | 20 (19%) | 20 (13%) | NA | 16 (18%) | 82 (36%) |
| High Likelihood | 48 (46%) | 68 (44%) | NA | 39 (44%) | 64 (28%) |
| Missing or unknown | 5 (5%) | 2 (1%) | NA | 5 (6%) | 0 |
| **Derived AD outcome†** |  |  |  |  |  |
| Control | 16 (15%) | 64 (41%) | 58 (18%) | 22 (25%) | 51 (22%) |
| AD | 66 (63%) | 77 (49%) | 180 (54%) | 52 (59%) | 125 (55%) |
| Other | 18 (17%) | 13 (8%) | 93 (28%) | 9 (10%) | 52 (23%) |
| Missing or unknown | 5 (5%) | 2 (1%) | 0 | 5 (6%) | 0 |

\*NIA Reagan score modified in accordance with Bennett et al, 2006 [63]: No AD: CERAD = No AD/None and Braak = Stage 0; Low Likelihood: CERAD = No AD/None and Braak ≥ Stage I OR CERAD = Possible/sparse and Braak = any stage OR CERAD = Probable AD/moderate and Braak ≤ Stage II; Intermediate Likelihood: CERAD = Probable/moderate and Braak ≥ Stage III OR CERAD = Definite/frequent and Braak ≥ Stage I and ≤ Stage IV; High Likelihood: CERAD = Definite AD/frequent and Braak ≥ Stage V.
†For Mayo patients, this outcome is the reported diagnosis according to Mayo neurologist guidelines, as reported [57]. For all other patients: Control: CERAD = No AD/None or Possible/sparse and Braak ≤ Stage III; AD case: CERAD = Probable/moderate or Definite/frequent and Braak ≥ Stage IV; Other = all other combinations of CERAD and Braak.
NA = not applicable for Mayo patients since Mayo did not report CERAD measures

### Sampling across brain regions

Different brain regions were sampled to capture differences in molecular profiles, including gene and protein expression across regions occurring at different stages of AD neuropathology (**Figure 1**). The dorsolateral prefrontal (DLPFC) cortex and temporal cortex are regions affected in AD, albeit typically later for DLPFC than the temporal cortex [69]. DLPFC [24] and temporal cortex--especially superior temporal gyrus (STG) [21,23]--were profiled with multi-omics measurements in AMP-AD studies of predominantly NHW donors. DLPFC and STG were obtained from all donors in the AMP-AD Diversity initiative, except those from Columbia, who had temporal pole tissue available instead of STG. The anterior caudate nucleus was selected as a non-cortical region also affected by AD neuropathology [70,71]. The total numbers of samples per tissue per data type and per donor by race and ethnicity are depicted in **Figure 2**. WGS for 626 donors were generated through the Diverse Cohorts initiative. It should be noted that WGS for an additional 408 donors for whom omics measures were generated in this study was readily available from the AMP-AD 1.0 initiative. Also, as mentioned earlier, Emory included samples from an additional 284 predominantly NHW donors from AMP-AD 1.0 to balance proteomics batches. The overlap between data contributor sites was generally highest for DLPFC and STG.

**Figure 1.**
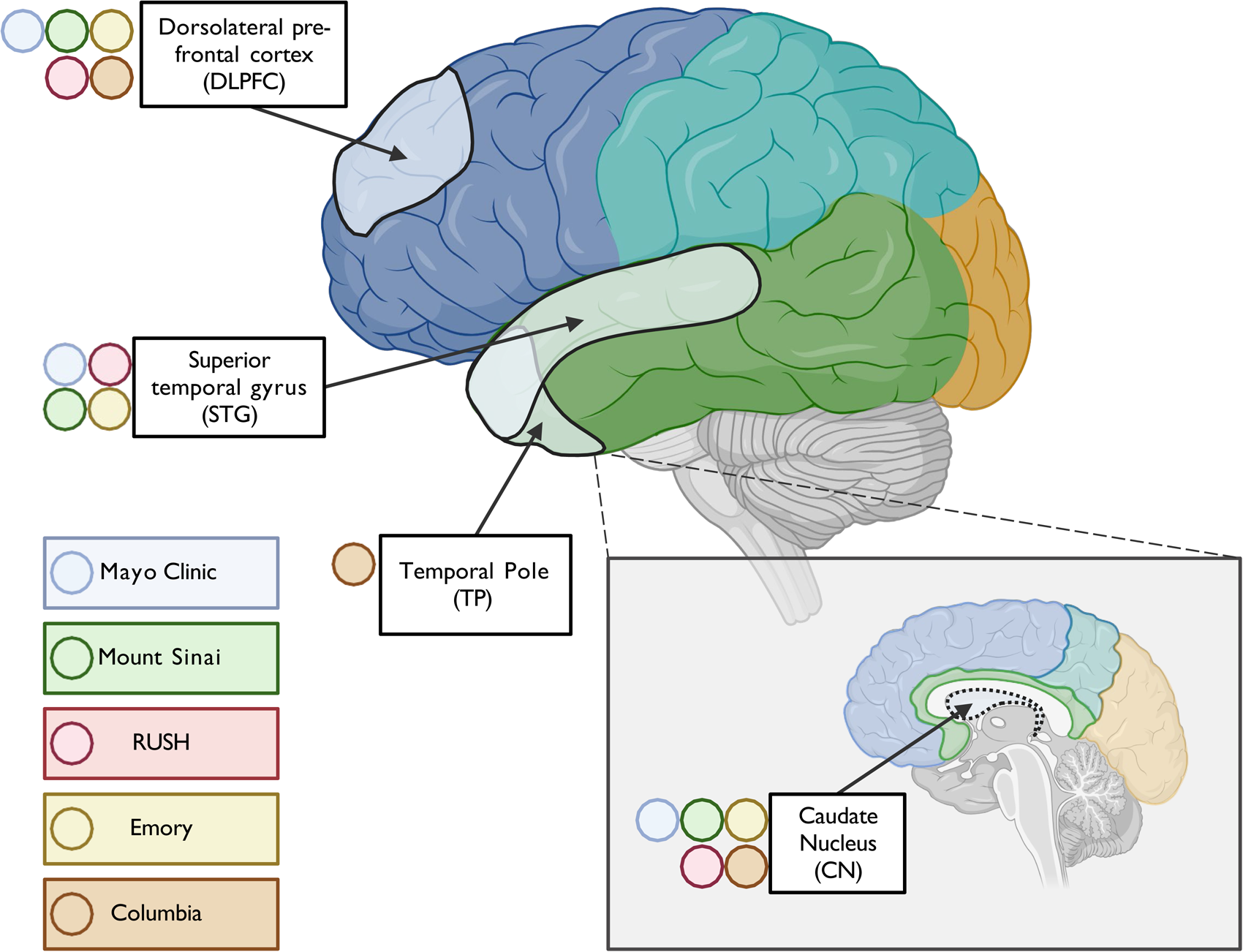
Profiled brain regions. Approximate location of tissue in brain regions sampled for molecular profiling, including RNAseq, WGS, and proteomics. Tissue from the dorsolateral prefrontal cortex (Brodmann areas 8, 9, and/or 46) and caudate nucleus were contributed by all sites, including Mayo Clinic, Mt. Sinai, Columbia, Rush, and Emory. In contrast, tissue from superior temporal gyrus (Brodmann 22) was provided by all sites except Columbia, which had only the temporal pole available for this lobe.

**Figure 2.**
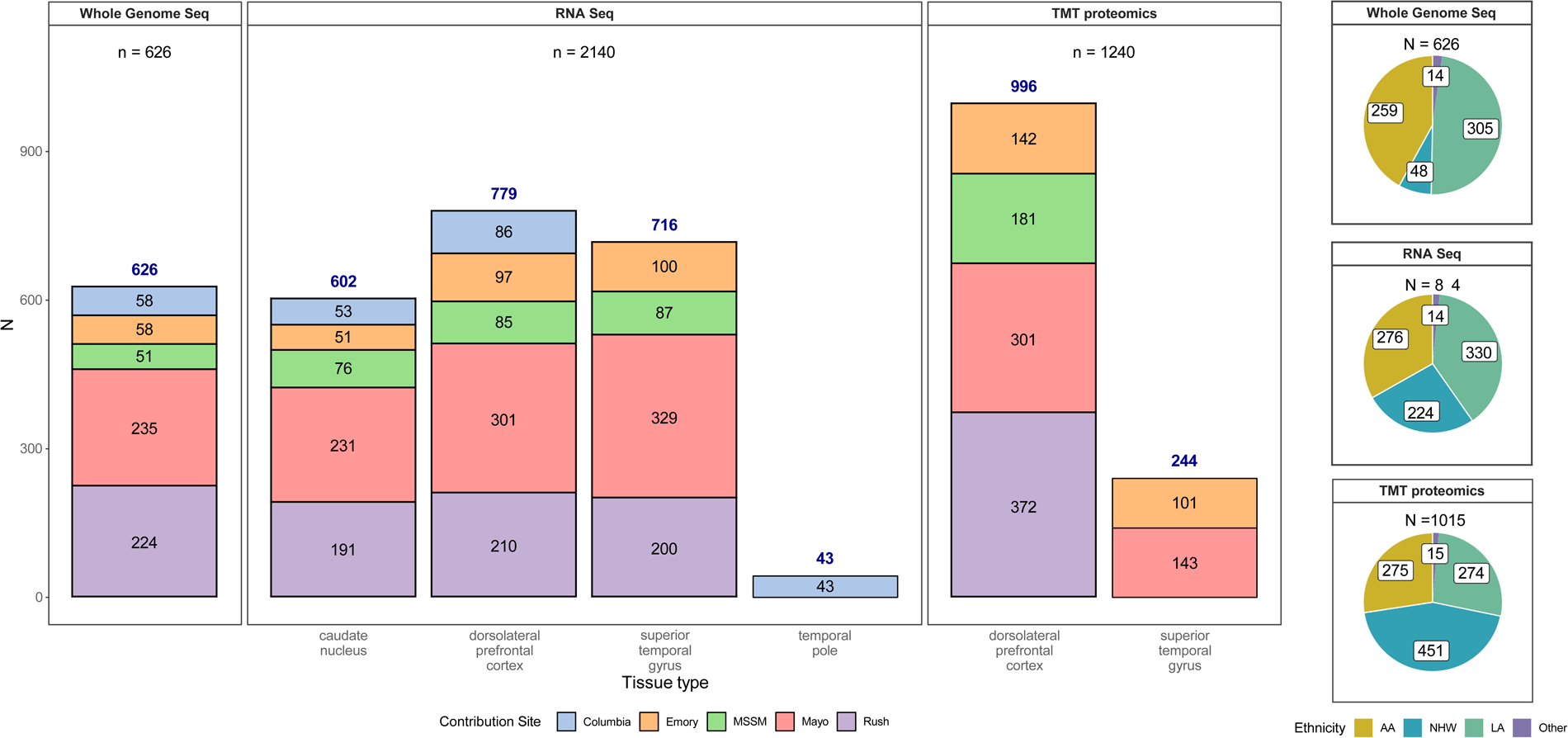
Data types by tissue, site, and individual race and ethnicity. Bar graph depicting the number of samples profiled by each assay (whole genome sequencing, RNAseq or TMT proteomics). Whole genome sequencing data was generated for 626 donors from various contributing sites (an additional 411 donors had WGS from AMP-AD 1.0 efforts, not shown here). Similarly, 2,140 unique transcriptomics profiles from RNAseq of caudate nucleus (n=602), dorsolateral prefrontal cortex (n=779), superior temporal gyrus (716) and temporal pole (n=43) from 844 donors were generated. Samples sent to other sites for the swap study are not included. A lone superior temporal gyrus RNAseq sample from Columbia was also not included in this summary. 1240 unique TMT-proteomes from dorsolateral prefrontal cortex (n=996) and superior temporal gyrus (n=244) were generated from 1,015 donors. These include the 284 samples from the AMP-AD 1.0 efforts to balance batches, as described in methods. Pie charts on the right show the number of donors profiled by ethnoracial categories (AA=African America, NHW=non-Hispanic White, LA=Latino American, and Other). These categories were defined as follows: donors whose race was encoded as “Black or African American” and ethnicity as ‘isHispanic=FALSE’ in the individual metadata were treated as ‘AA’. Those with race encoded as White and ethnicity as ‘isHispanic=FALSE’ were categorized as ‘NHW’. Remaining donors, for whom ethnicity was encoded as ‘isHispanic=TRUE’ were treated as ‘LA’. All remaining donors from various other races were encoded as ‘Other’.

### DNA Extraction

All DNA extractions were done from the dorsolateral prefrontal cortex for subsequent whole genome sequencing (WGS). Mayo Clinic extracted DNA for all samples from the Mayo Clinic, Banner Sun Health, University of Florida, and Emory University Brain Banks. DNA was manually extracted from frozen brain tissue and was isolated using the AutoGen245T Reagent Kit (Part #agkt245td) according to the manufacturer’s protocol, including an Rnase step (Qiagen, Cat# 19101) following tissue digestion. DNA was quantified for amount and purity using the Nanodrop Spectrophotometer (ThermoFisher, Waltham, MA) and Qubit 2.0 Fluorometer (ThermoFisher, Waltham, MA). 1875 ng per donor were transferred on dry ice to the New York Genome Center (NYGC) for whole genome library preparation and sequencing (WGS). For all other samples, DNA extraction was performed at the NYGC. In brief, for Rush and Mount Sinai samples, 25 mg of tissue was homogenized using a Qiagen Buffer ATL/Proteinase K with overnight incubation at 56 degrees Celsius. DNA was extracted using the Qiagen QIAamp DNA Mini Kit (Qiagen, 51304), and a Qiagen QIAamp DNA Mini Kit (Qiagen, 51304) was used for DNA cleanup. For Columbia samples, 50 mg of tissue was homogenized using a Buffer TE/Rnase A Solution (Maxwell Cat.# A7973). DNA was extracted using a Promega Maxwell kit (AS1610) and cleaned using a Maxwell RSC Tissue DNA Kit (Maxwell, TM476). For all samples, DNA quality was analyzed using a Fragment Analyzer (Advanced Analytics) or BioAnalyzer (Agilent Technologies). Libraries were generated using the Illumina Tru-Seq PCR-Free protocol, and WGS was performed by the NYGC.

### Whole Genome Sequencing (WGS)

NYGC performed QC on the raw WGS reads and provided the following metrics: Total Reads, PF Reads, % PF Reads, PF Aligned Reads, % PF Aligned, PF Aligned Pairs, % PF Aligned Pairs, Mean Read Length, Strand Balance, Estimated Library Size, Mean Coverage, % Sequence Contamination, Median Insert Size, Mean Insert Size, AT Dropout, GC Dropout, and % Total Duplication. These metrics were generated using Picard tools (v2.4.1, http://picard.sourceforge.net) following paired-end read alignment to the GRCh38 human reference using the Burrows-Wheeler Aligner (BWA-MEM v0.7.15). Sequence contamination was estimated on a per-sample basis using VerifyBamID (https://genome.sph.umich.edu/wiki/VerifyBamID).

### RNA extraction

RNA extractions were done from frozen tissue from the dorsolateral prefrontal cortex, anterior caudate nucleus, and superior temporal gyrus (or temporal pole) (**Figure 1**) for subsequent RNA sequencing. Most donors had tissue from all 3 regions, but no donors were excluded for lacking samples from any brain regions. Brain tissue from Emory, Banner, and the University of Florida was sent to Mayo Clinic Jacksonville in Florida for RNA isolation and sequencing. Brain tissue samples for the Mayo cohort were obtained from the Mayo Clinic Brain Bank. RNA was isolated using a Trizol/chloroform protocol, followed by 2-step RNA purification (Qiagen Rneasy Mini Kit) and concentration incorporating on-column (Qiagen Cat#74106 or 74104 and Cat#79254) and liquid (Zymo Cat# R1014 or R1013) Dnase steps respectively. The quantity and quality of all RNA samples were determined by the NanoDrop 2000 Spectrophotometer and Agilent 2100 Bioanalyzer using the Agilent RNA 6000 Nano Chip (Cat# 5067-1511 from Agilent Technologies, Santa Clara, CA).

For all Rush samples, 50mg of frozen brain tissue was dissected and homogenized in DNA/RNA shield buffer (Zymo, R1100) with 3mm beads using a bead homogenizer. RNA was subsequently extracted using Chemagic RNA tissue kit (Perkin Elmer, CMG-1212) on a Chemagic 360 instrument. RNA was concentrated (Zymo, R1080), and RQN values were calculated with a Fragment Analyzer total RNA assay (Agilent, DNF-471).

Tissue samples from MSSM and Columbia were prepared for RNA sequencing at the NYGC. Tissue was homogenized using TRIzol (needles), and RNA was extracted using Cloroform. A Qiagen Rneasy Mini Kit was used for RNA cleanup, and quality was analyzed with Fragment Analyzer (Advanced Analytics) or BioAnalyzer (Agilent Technologies).

### RNA sequencing

Brain samples from the Mayo Clinic, Banner Sun Health, University of Florida, and Emory University were randomized with respect to race, ethnicity, diagnosis (AD, control, other), contributing institution, RIN, *APOE* genotypes, sex, and age prior to transfer to the Mayo Clinic Genome Analysis Core for library preparation and sequencing across 13 flowcells. Total RNA concentration and quality were determined using Qubit fluorometry (ThermoFisher Scientific, Waltham, MA) and the Agilent Fragment Analyzer (Santa Clara, CA). Using Illumina’s TruSeq Stranded Total RNA reagent kit [Cat #20020597] and the Illumina Ribo-Zero Plus rRNA Depletion kit [Cat #20037135] (San Diego, CA), libraries were prepared according to the manufacturer’s instructions with 200 ng of total RNA. The concentration and size distribution of the completed libraries were determined using Qubit fluorometry and the Agilent TapeStation D1000 (Santa Clara, CA). Libraries were sequenced at an average of 200M total reads, following the standard protocol for the Illumina NovaSeq 6000. The flow cell was sequenced as 100 X 2 paired-end reads using the NovaSeq S4 sequencing kit and NovaSeq Control Software v1.7.5. Base-calling was performed using Illumina’s RTA version 3.4.4. All RNA samples isolated from tissue samples of the same donor were sequenced together in the same flowcell.

For all Rush samples, following RNA extraction, concentration was determined using Qubit broad-range RNA assay (Invitrogen, Q10211) according to the manufacturer’s instructions. 500ng total RNA was used as input for sequencing library generation, and rRNA was depleted with RiboGold (Illumina, 20020599). A Zephyr G3 NGS workstation (Perkin Elmer) was utilized to generate TruSeq stranded sequencing libraries (Illumina, 20020599) with custom unique dual indexes (IDT) according to the manufacturer’s instructions with the following modifications. RNA was fragmented for 4 minutes at 85°C. The first strand synthesis was extended to 50 minutes. Size selection post adapter ligation was modified to select for larger fragments. Library size and concentrations were determined using an NGS fragment assay (Agilent, DNF-473) and Qubit ds DNA assay (Invitrogen, Q10211), respectively, according to the manufacturer’s instructions. The modified protocol yielded libraries with an average insert size of around 330-370bp. Libraries were normalized for molarity and sequenced on a NovaSeq 6000 (Illumina) at 40-50M reads, 2x150bp paired end.

Columbia and MSSM samples were sequenced at the NYGC. Following rRNA depletion using RiboErase, libraries were prepared using 500 ng of RNA with the KAPA Stranded Total RNA (HMR) RiboErase Kit (kapabiosystems). RNA was fragmented for 5 minutes at 85°C, and first strand synthesis was extended to 10 min at 25°C, 15 min at 42°C, and 15 min at 70°C. Size selection post adapter ligation was modified to select larger fragments, which resulted in 480-550 bp fragments. Sequencing was performed using an Illumina NovaSeq 6000 to generate 100bp paired-end reads. Sequencing quality control was performed using Picard version 1.83 and RseQC version 2.6.1. STAR version 2.5.2a was used to align reads to the GRCh38 genome using Gencode v25 annotation. Bowtie2 version 2.1.0 was used to measure rRNA abundance. Annotated genes were quantified with featureCounts version 1.4.3-p1. Sequence contamination was estimated on a per-sample basis using VerifyBamID (https://genome.sph.umich.edu/wiki/VerifyBamID). The identity of the RNA sample is confirmed by evaluating concordance with whole genome sequencing data using Conpair, a tool that uses a set of SNPs common in the human population to determine sample identity.

To maximize the number of brain samples included in the AMP-AD Diversity Initiative, RNA Integrity Number (RIN) was measured but not used to filter out samples. The DV200 is an assessment of the proportion of RNA fragments greater than 200 nucleotides and is considered a more accurate measure of RNA quality when RIN value is low [72]. For Columbia and RUSH, at least 85% of the low RIN samples (RIN<5) have a DV200 > 70%; for Mayo, 90% of samples meet this metric, and for MSSM, 95% of the sample pass. Given the high proportion of samples with a DV200 >70%, samples were not removed based on these metrics but rather assessed carefully at the QC stage.

### RNA sample exchange

Since RNA sequencing was conducted at three different sequencing centers (Mayo Genome Analysis Core, NYGC, and Rush), a small number of samples were exchanged between the three sequencing centers to evaluate the extent of technical variability between these centers (**Figure 3**). Mayo Clinic contributed 5 samples each from the dorsolateral prefrontal cortex (DLPFC) and superior temporal gyrus to Rush and NYGC. 6 DLPFC samples from Columbia were sent to Mayo Clinic and Rush, and 4 samples each for DLPFC and STG from Mt. Sinai were sent to Mayo and Rush. Rush contributed 6 samples each from DLPFC and STG to the Mayo Clinic and NYGC. Tissues sent to other sites as part of the swap experiment were also sequenced at each original sequencing site, resulting in 3 sets of RNAseq data from each participant and brain region for the swapped samples. RNA extraction and sequencing protocol for swap samples at each site is described above (see RNA extraction and RNA sequencing).

**Figure 3.**
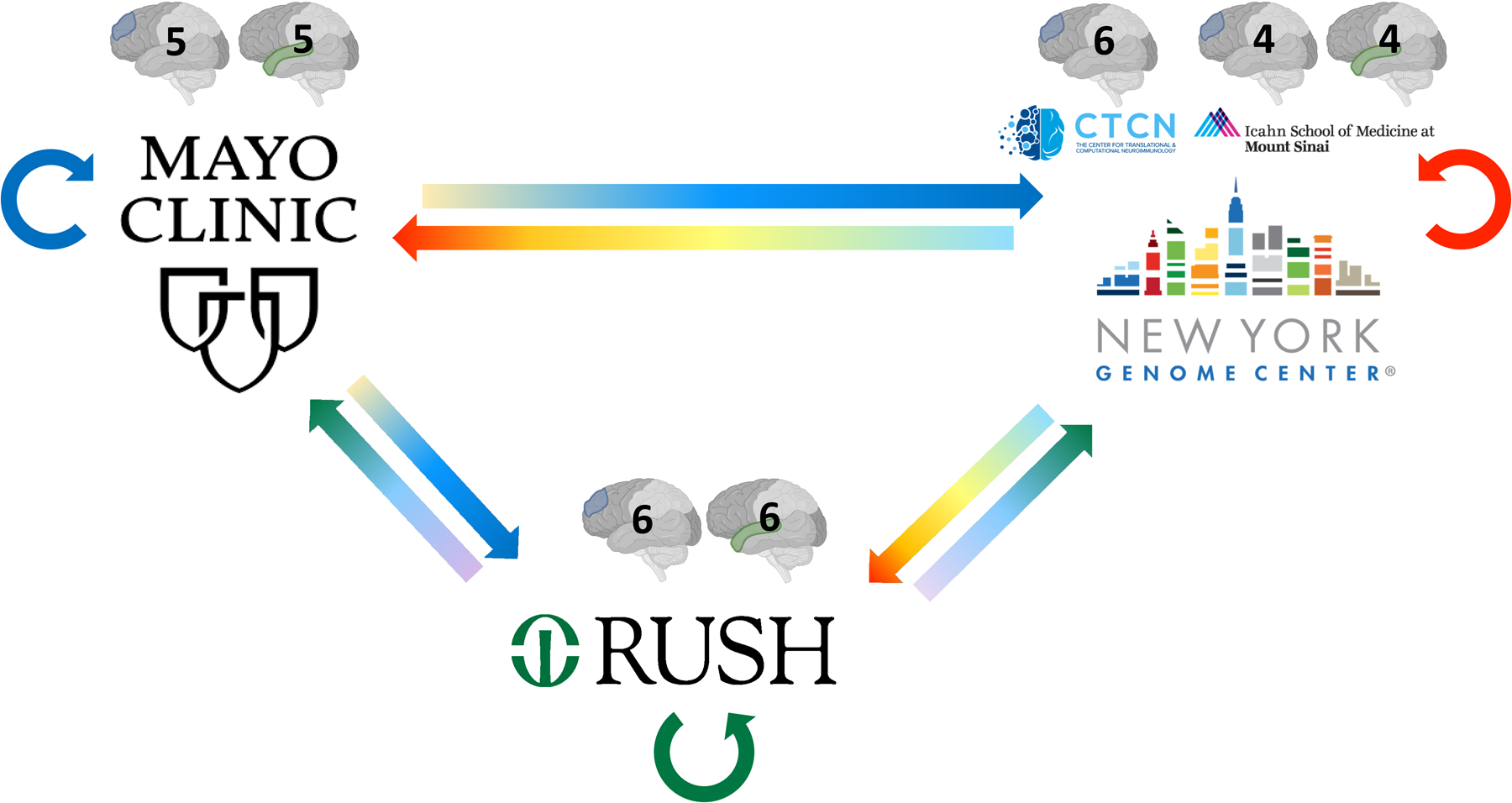
RNAseq sample swaps. To evaluate the technical variability of RNA sequencing amongst the three sites, RNA tissue from the same brain was sequenced at each site for a small number of samples. The number and region of samples exchanged are illustrated with the grayscale brain image with the exchanged tissue highlighted in color (DLPFC in blue, STG in green). Straight arrows represent tissue exchange; circular arrows represent tissue sequenced at the original site, shown in blue, green, and red circular arrows for Mayo Clinic, Rush, and NYGC, respectively. Samples from MSSM (4 DLPFC, 4 STG) and Columbia (5 DLPFC) were utilized for the swap experiment at NYGC.

All samples that were part of the swap study were sequenced in a single batch at Mayo, whereas samples sequenced at NYGC were distributed across 5 batches, and at Rush, they were distributed across 3 batches. RIN values for samples sequenced at Mayo ranged between 2.7 and 8.8, whereas those at NYGC ranged between 2.7 and 8.7, and at Rush ranged between 1.3 and 8.0. RNAseq data for swap samples generated across all three sites were consensus processed using MAPRSeq v3 pipeline [73]. Reads were aligned to the reference (GRCh38) using STAR aligner v2.6.1. Sequencing and alignment metrics from FastQC and RseQC were utilized to evaluate variability across sequencing centers. The median base quality of reads was consistent (Phred ≥ 37) across sites for both DLPFC and STG. Evaluation of base content (percentage of As, Ts, Gs, and Cs at each position in the read) between the 25th and 75th percentile along the read length revealed that the percentage of As and Ts was around 30% and that of Gs and Cs was 20% across all reads and samples. The following summary metrics are summarized by tissue contribution site and sequencing site in **Supplementary** Figure 1. Between 104 and 147 million (M) reads were generated for samples sequenced at Mayo, 95 to 98% of which were mapped to the genome and 31 to 54% mapped to genes. For samples sequenced at NYGC, between 58 and 222M reads were generated, 93 to 98% of which mapped to the genome and 37 to 58% mapped to genes. Similarly, at Rush, between 10 and 125M reads were generated, 83 to 96% mapped to the genome and 28 to 57% mapped to genes. The median ratio of reads covering the 80^th^ and 20^th^ percentile along the gene body for all genes was between 1 and 1.1, revealing no significant bias towards 3’ or 5’ degradation. Sex deduced from gene expression was consistent with assigned sex based on clinical information. After conditional quantile normalization (CQN) to identify expressed genes, principal component analysis (PCA) was performed to evaluate stratification amongst samples (**Supplementary** Figure 2). When PCs were generated by tissue (one set of PCs each of DLPFC and STG) and plotted together, there was no separation by tissue contribution site (**Supplementary** Figure 2a), although there was some separation by sequencing site (**Supplementary** Figure 2b), and indeed, sequencing site was the largest source of technical variation. When PCs were generated by tissue contribution site (one set of PCs each for Columbia, Mt. Sinai, Mayo, and Rush) and plotted together, there was no separation by sequencing site but only by tissue (**Supplementary** Figure 2c).

### Proteomics

Proteome measurements were conducted in all DLPFC tissue, as well as in STG, for a subset of the samples from the Mayo Clinic to enable joint analyses with other STG proteome data from this Brain Bank [21]. Pre-and post-processing steps for proteomic quantification were performed at Emory University for all samples from all contributing institutions using the following methods. Samples from each individual site were randomized in batches of 15 to 17 and balanced, where possible, with respect to race, ethnicity, diagnosis (AD), sex, age, [74]. Batching schema is included in the proteomics biospecimen metadata file (syn53185805).

#### Brain tissue homogenization and protein digestion

Procedures for tissue homogenization for all tissues were performed essentially as described [48,75]. Approximately 100 mg (wet tissue weight) of brain tissue was homogenized in 8 M urea lysis buffer (8 M urea, 10 mM Tris, 100 mM NaHPO4, pH 8.5) with HALT protease and phosphatase inhibitor cocktail (ThermoFisher) using a Bullet Blender (NextAdvance) essentially as described [75]. Each Rino sample tube (NextAdvance) was supplemented with ∼100 μL of stainless steel beads (0.9 to 2.0 mm blend, NextAdvance) and 500LJμL of lysis buffer. Tissues were added immediately after excision, and samples were placed into the bullet blender at 4 °C. The samples were homogenized for 2 full 5 min cycles, and the lysates were transferred to new Eppendorf Lobind tubes. Each sample was then sonicated for 3 cycles of 5 s of active sonication at 30% amplitude, followed by 15 s on ice. Samples were centrifuged for 5 min at 15,000LJx g, and the supernatant was transferred to a new tube. Protein concentration was determined by bicinchoninic acid (BCA) assay (Pierce). For protein digestion, 100LJμg of each sample was aliquoted, and volumes were normalized with additional lysis buffer. An equal amount of protein from each sample was aliquoted and digested in parallel to serve as the global pooled internal standard (GIS) in each TMT batch, as described below. Similarly, GIS pooled standards were generated from all cohorts. Samples were reduced with 1LJmM dithiothreitol (DTT) at room temperature for 30 min, followed by 5LJmM iodoacetamide (IAA) alkylation in the dark for another 30 min. Lysyl endopeptidase (Wako) at 1:100 (w/w) was added, and digestion was allowed to proceed overnight. Samples were then 7-fold diluted with 50 mM ammonium bicarbonate. Trypsin (Promega) was added at 1:50 (w/w), and digestion was carried out for another 16 h. The peptide solutions were acidified to a final concentration of 1% (vol/vol) formic acid (FA) and 0.1% (vol/vol) trifluoroacetic acid (TFA), and desalted with a 30 mg HLB column (Oasis). Each HLB column was first rinsed with 1LJmL of methanol, washed with 1 mL 50% (vol/vol) acetonitrile (ACN), and equilibrated with 2×1 mL 0.1% (vol/vol) TFA. The samples were loaded onto the column and washed with 2×1 mL 0.1% (vol/vol) TFA. Elution was performed with 2 volumes of 0.5 mL 50% (vol/vol) ACN. The eluates were then dried to completeness using a SpeedVac.

#### Isobaric Tandem Mass Tag (TMT) Peptide Labeling

The Synapse DOI giving sample to batch arrangement is presented Table 4. In preparation for labeling, each brain peptide digest was resuspended in 75 μl of 100 mM triethylammonium bicarbonate (TEAB) buffer; meanwhile, 5 mg of TMT reagent was dissolved into 200 μl of ACN. Each sample (containing 100 μg of peptides) was re-suspended in 100LJmM TEAB buffer (100 μL). The TMT labeling reagents (5mg; Tandem Mass Tag (TMTpro) kit (Thermo Fisher Scientific, A44520)) were equilibrated to room temperature, and anhydrous ACN (256 μL) was added to each reagent channel. Each channel was gently vortexed for 5 min, and then 41 μL from each TMT channel was transferred to the peptide solutions and allowed to incubate for 1 h at room temperature. The reaction was quenched with 5% (vol/vol) hydroxylamine (8 μl) (Pierce). All channels were then combined and dried by SpeedVac (LabConco) to approximately 150 μL and diluted with 1 mL of 0.1% (vol/vol) TFA, then acidified to a final concentration of 1% (vol/vol) FA and 0.1% (vol/vol) TFA. Labeled peptides were desalted with a 200 mg C18 Sep-Pak column (Waters). Each Sep-Pak column was activated with 3 mL of methanol, washed with 3 mL of 50% (vol/vol) ACN, and equilibrated with 2×3 mL of 0.1% TFA. The samples were then loaded and each column was washed with 2×3 mL 0.1% (vol/vol) TFA, followed by 2 mL of 1% (vol/vol) FA. Elution was performed with 2 volumes of 1.5 mL 50% (vol/vol) ACN. The eluates were then dried to completeness using a SpeedVac.

**Table 4.**
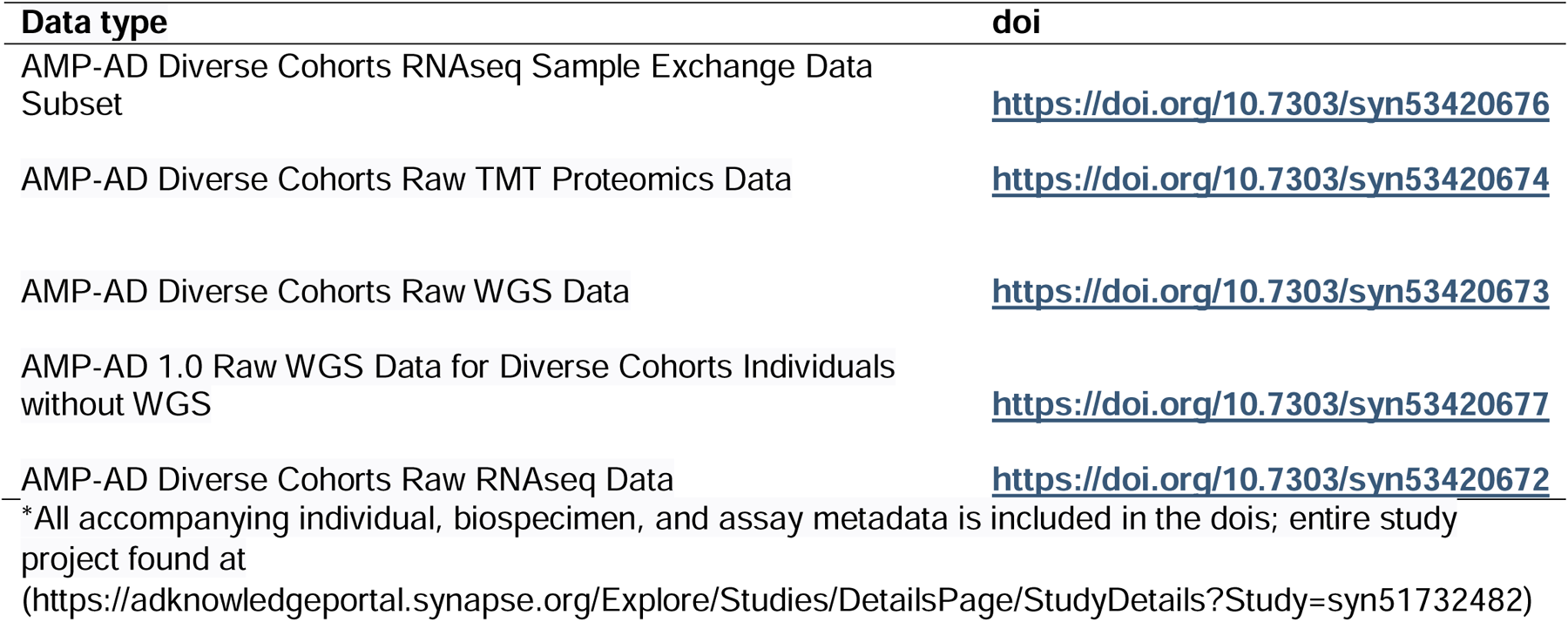
Synapse doi’s of data shared on the AD Knowledge Portal for the AMP-AD Diversity Initiative*.

| **Data type** | **doi** |
| --- | --- |
| AMP-AD Diverse Cohorts RNAseq Sample Exchange Data Subset | https://doi.org/10.7303/syn53420676 |
| AMP-AD Diverse Cohorts Raw TMT Proteomics Data | https://doi.org/10.7303/syn53420674 |
| AMP-AD Diverse Cohorts Raw WGS Data | https://doi.org/10.7303/syn53420673 |
| AMP-AD 1.0 Raw WGS Data for Diverse Cohorts Individuals without WGS | https://doi.org/10.7303/syn53420677 |
| AMP-AD Diverse Cohorts Raw RNAseq Data | https://doi.org/10.7303/syn53420672 |

\*All accompanying individual, biospecimen, and assay metadata is included in the dois; entire study project found at (https://adknowledgeportal.synapse.org/Explore/Studies/DetailsPage/StudyDetails?Study=syn51732482)

#### High-pH off-line fractionation

High-pH fractionation was performed essentially as described with slight modification [75,76]. Dried samples were re-suspended in high pH loading buffer (0.07% vol/vol NH_4_OH, 0.045% vol/vol FA, 2% vol/vol ACN) and loaded onto a Water’s BEH 1.7 um 2.1mm by 150mm. A Thermo Vanquish or Agilent 1100 HPLC system was used to carry out the fractionation. Solvent A consisted of 0.0175% (vol/vol) NH_4_OH, 0.01125% (vol/vol) FA, and 2% (vol/vol) ACN; solvent B consisted of 0.0175% (vol/vol) NH_4_OH, 0.01125% (vol/vol) FA, and 90% (vol/vol) ACN. The sample elution was performed over a 25 min gradient with a flow rate of 0.6 mL/min. A total of 192 individual equal volume fractions were collected across the gradient and subsequently pooled by concatenation into 96 fractions (RUSH, MSSB, and Mayo cohorts) or 48 fractions for the Emory Cohort. All peptide fractions were dried to completeness using a SpeedVac. Off-line fractionation of the Mount Sinai and Emory cohorts was performed as previously described [75,77].

#### TMT mass spectrometry

All fractions were resuspended in an equal volume of loading buffer (0.1% FA, 0.03% TFA1% ACN) and analyzed by liquid chromatography coupled to tandem mass spectrometry essentially as described [78], with slight modifications. Peptide eluents were separated on a self-packed C18 (1.9 μm, Dr. Maisch, Germany) fused silica column (25 cm × 75 μM internal diameter (ID); New Objective, Woburn, MA) by a Dionex UltiMate 3000 RSLCnano liquid chromatography system (ThermoFisher Scientific) and monitored on a mass spectrometer (ThermoFisher Scientific). Sample elution was performed over a 180 min gradient with a flow rate of 225 nL/min. The gradient was from 3% to 7% buffer B over 5 min, then 7% to 30% over 140 min, then 30% to 60% over 5 min, then 60% to 99% over 2 min, then held constant at 99% solvent B for 8 min, and then back to 1% B for an additional 20 min to equilibrate the column. The mass spectrometer was set to acquire data in data-dependent mode using the top-speed workflow with a cycle time of 3 seconds. Each cycle consisted of 1 full scan followed by as many MS/MS (MS2) scans that could fit within the time window. The full scan (MS1) was performed with an m/z range of 350-1500 at 120,000 resolution (at 200 m/z) with AGC set at 4x10^5^ and a maximum injection time of 50 ms. The most intense ions were selected for higher energy collision-induced dissociation (HCD) at 38% collision energy with an isolation of 0.7 m/z, a resolution of 30,000, an AGC setting of 5x10∧4, and a maximum injection time of 100 ms. Of the 72 TMT batches for the dorsolateral pre-frontal cortex tissues, 34 were run on an Orbitrap Fusion Lumos mass spectrometer, 24 batches were run on an Orbitrap Fusion Eclipse GC 240 mass spectrometer, and 14 batches were run on an Orbitrap Eclipse mass spectrometer as previously described [75]. Collectively, LC-MS/MS led to a total of 6479 raw files from frontal cortex, and 1824 raw files from temporal cortex tissue samples (Fig. 1A), with the distribution as follows: Emory University Frontal Cortex Cohort: 431; Mayo Clinic Frontal Cortex Cohort: 2304; Mount Sinai Frontal Cortex Cohort: 1344; Rush University Frontal Cortex Cohort: 2400; and Emory University and Mayo Clinic Temporal Cortex Cohort: 1824.

## Discussion

This is a data descriptor study for the AMP-AD [20] Diversity Initiative that was launched to generate, analyze, and make available to the research community multi-omics data in AD and older control brain donors from multi-ethnic populations enriched for AA and LA participants who are at higher risk [2] for AD but traditionally underrepresented in research [3–6]. While GWAS in AA and LA participants are orders of magnitude smaller than that for NHW, multi-omics studies are essentially non-existent, especially in brain tissue from these populations. This underrepresentation in brain multi-omics studies is in part due to lower autopsy rates in AA and LA populations [79,80], the causes of which are multi-factorial but must be comprehensively understood to overcome this barrier in research. There are efforts to increase diversity in autopsy studies for ADRD [63,81,82], which have led to the discovery that some but not all neuropathologies have ethnoracial differences [81,83–85].

To our knowledge, there are no sizable multi-omics studies of ADRD including age-matched control AA and LA donors to uncover the molecular underpinnings of these neuropathologies. In contrast, the AMP-AD Target Discovery and Preclinical Validation Project generated [21–24] and broadly shared [25] multi-omics data on >2,500 brain samples, primarily from NHW donors. These multi-omics data revealed brain molecular alterations in specific biological pathways, including but not limited to innate immunity, synaptic biology, myelination, vascular biology, and mitochondrial energetics [28–30,32–34,37,39,45,54,86–89], thereby supporting complex, heterogeneous molecular etiologies, resulting in >600 therapeutic candidates with a step closer to precision medicine in ADRD.

Recognizing the essential importance of inclusivity in precision medicine [56], we launched the AMP-AD Diversity Initiative with the objective of performing multi-omics profiling and analysis of samples from diverse cohorts to discover the full spectrum of therapeutic targets and biomarkers that will be of utility to all populations affected with AD. In this data descriptor manuscript, we describe the first wave of data generated and shared with the research community, comprising transcriptome from three brain regions, whole genome sequence, and proteome measures from 908 multi-ethnic donors enriched for AA (n=306) and LA (n=326). We emphasize that this is the initial set of data currently being expanded to include other omics measures, namely metabolome, single-cell RNAseq, and epigenome in the AMP-AD Diverse Cohorts Study.

We must emphasize that multi-omics studies alone are unlikely to be sufficient to discover all causes of ADRD or explain the disparities in risk observed for AA and LA participants [4,6,90]. Rather, this requires a full understanding of the role of the exposome, including sex, race, ethnicity, lifetime health measures, co-morbidities, and additional structural and social determinants of health (SSDoH) [54,91–96]. Only by capturing the exposome and evaluating its complex interactions with multi-omics measures and disease-related outcomes can we have a holistic lens into the etiopathogenesis of ADRD. With this goal in mind, the AMP-AD Diversity Initiative is in the process of curating and harmonizing exposome data for the donors in the AMP-AD Diverse Cohorts Study.

Despite the potential utility of this foundational multi-omics dataset from a multi-ethnic autopsy cohort, there are shortcomings in the current study. To include the largest possible number of AA and LA donors, brain tissue from both archival brain banks and longitudinal studies was included, resulting in variability in the types of clinical and neuropathologic data available. We strove to overcome this variability by careful harmonization of the neuropathologic data to the extent possible, although must underscore the need to have more diverse autopsy cohorts with in-depth and uniform phenotyping, including clinical and neuropathologic variables. For this study, we accepted self-reported race and ethnicity. We recognize that race and ethnicity are highly complex constructs [6,80,90,97] that must consider SSDoH, cultural, historical, and biological variables and context. While we will aim to incorporate as many exposome variables into this study as possible, there is clearly a need for multi-disciplinary teams to assess all non-biological and biological variables and context holistically in large-scale population-based studies to understand disparities in and causes of disease risk. Finally, though our study is a step in the right direction for inclusivity in precision medicine studies, there are many other underrepresented groups in ADRD research in the United States and globally [3,79]. National and global initiatives are required to expand this research to all affected populations.

In summary, we describe transcriptome data from 2224 brain samples, proteome data from 1385 samples, and new whole genome sequencing from 626 samples, primarily from 908 multi-ethnic donors enriched for AA and LA participants. This data is accompanied by harmonized neuropathologic diagnoses of AD (n=500), control (n=211), or other (n=185). These data made available to the research community are expected to be an initial step to bridge our data and knowledge gap in the understanding of AD in underrepresented and -at-risk populations.

## Data Availability

The data described herein is available for use by the research community and has been deposited in the AD Knowledge Portal, with all publicly available data found under The Accelerating Medicines Partnership Alzheimer’s Disease Diverse Cohorts Study (AMP-AD Diverse Cohorts Study): (https://adknowledgeportal.synapse.org/Explore/Studies/DetailsPage/StudyDetails?Study=syn51732482). **Table 4** provides a list of the files and folders containing all data, their specific Synapse identifiers (IDs), DOIs, and brief descriptions of the file or folder contents. These files and their assigned DOIs will be maintained in perpetuity in the AMP-AD Knowledge Portal. Access to all of these files is enabled through the Sage Bionetworks, Synapse repository.

The AD Knowledge Portal hosts data from multiple cohorts that were generated as part of or used in support of the AMP-AD Diverse Cohorts Study conducted under the AMP-AD Diversity Initiative. The portal uses the Synapse software platform for backend support, providing users with web-based and programmatic access to data files. All data files in the portal are annotated using a standard vocabulary to enable users to search for relevant content across the AMP-AD datasets using programmatic queries. Data is stored in a cloud-based manner hosted by Amazon web services (AWS), which enables users to execute cloud-based compute or copy the data to local infrastructure. Detailed descriptions, including data processing, QC metrics, and assay and cohort-specific variables, are provided for each file as applicable.

Access to the data described herein is controlled in a manner set forth by the institutional review boards (IRB) at the Mayo Clinic, MSSM, Rush, Emory, and Columbia. All data use terms include (1) maintenance of data in a secure and confidential manner, (2) respect for the privacy of study participants, (3) including the following in any published text: “The results published here are in whole or in part based on data obtained from the AD Knowledge Portal (https://adknowledgeportal.org/). Data generation was supported by the following NIH grants: U01AG046139, U01AG046170, U01AG061357, U01AG061356, U01AG061359, and R01AG067025. We thank the participants of participants of the Religious Order Study, Memory and Aging Project, the Minority Aging Research Study, Rush Alzheimer’s Disease Research Center, Mount Sinai/JJ Peters VA Medical Center NIH Brain and Tissue Repository, National Institute of Mental Health Human Brain Collection Core (NIMH HBCC), Mayo Clinic Brain Bank, Sun Health Research Institute Brain and Body Donation Program, Goizueta Alzheimer’s Disease Research Center, New York Brain Bank at Columbia University, New York Genome Center and the Biggs Institute Brain Bank for their generous donations. Data and analysis contributing investigators include Nilüfer Ertekin-Taner, Minerva Carrasquillo, Mariet Allen, Dennis Dickson (Mayo Clinic, Jacksonville, FL), David Bennett, Lisa Barnes (Rush University), Philip De Jager, Vilas Menon (Columbia University), Bin Zhang, Vahram Haroutanian (Icahn School of Medicine at Mount Sinai), Allan Levey, Nick Seyfried (Emory University), Rima Kaddurah-Daouk (Duke University), Steve Finkbeiner (University of California-San Francisco/Gladstone Institutes), Daifeng Wang (University of Wisconsin-Madison), Stefano Marenco (NIMH HBCC), Anna Greenwood, Abby Vander Linden, Laura Heath, William Poehlman (Sage Bionetworks).” For access to content described in this manuscript see: https://doi.org/10.7303/syn53420672, https://doi.org/10.7303/syn53420673, https://doi.org/10.7303/syn53420674, https://doi.org/10.7303/syn53420676, https://doi.org/10.7303/syn53420677 (also listed in **Table 4**). To download data, users must register for a Synapse account, provide electronic agreement to the Terms of Use outlined above, and complete a Data Use Certificate. User approvals are managed by the Synapse Access and Compliance Team (ACT).

## Author Contributions

J.S.R, L.H., N.E-T wrote the initial draft of the manuscript. J.S.R., L.H., A.V.L., M.A., A.G., N.E-T. collated and oversaw the organization of data and samples for the AMP-AD Diversity Initiative. J.S.R, M.A., K.d.P.L., E.J.F, E.W., Y.M., S.P, T.B., A.T, V.H., M.G, D.W.D., M.G., and E.B.L. provided and organized brain samples from the Mayo Clinic, Rush, Emory, Upenn, Mount Sinai, Columbia, Banner and the University of Florida Brain Banks. F.S., L.Y., K.X., L.P., E.S.M., E.B.D., A.S., L.P., Z.Q, .J.S.R., E.J.F., A.P.W., T.S.W., W.P., Z.Q., A.R., Y.W., D.M.D., E.M., and S.R.O. analyzed the transcriptome, genome, and proteome data. L.H., A.V.L., M.A., J.S., C.H., M.M.C, M.Atik., G.Y., A.M., T.T.N., S.P, T.B., A.T, V.H., M.G, and D.W.D. provided data and performed analyses for phenotype harmonization. H.R., H.X., S.P, T.B., A.T, V.H., M.G, and D.W.D. provided neuropathology measures. S.S., R.M., L.B., P.D.J., B.Z., D.B., J.J.L., A.I.L., D.X.M., N.S., and N.E-T. led the cohort studies from which donor tissue and data are obtained. P.D.J., B.Z., D.B., N.S., A.G., and N.E-T. obtained funding for and designed the AMP-AD Diversity Initiative and provided supervision. All authors reviewed and provided feedback for the manuscript.

## Acknowledgements and Funding Sources

We would like to thank the patients and their families for their participation; without them, these studies would not have been possible. The results published here are based on data available in the AD Knowledge Portal (https://adknowledgeportal.org). The Mayo RNAseq study data was led by Dr. Nilüfer Ertekin-Taner, Mayo Clinic, Jacksonville, FL, as part of the multi-PI U01 AG046139 (MPIs Golde, Ertekin-Taner, Younkin, Price) using samples from The Mayo Clinic Brain Bank. Data collection was supported through funding by NIA grants P50 AG016574, R01 AG032990, U01 AG046139, R01 AG018023, U01 AG006576, U01 AG006786, R01 AG025711, R01 AG017216, R01 AG003949, P30AG072979, P01AG066597, U19AG062418, U01AG061357, RF1AG062181, P30AG066511 CurePSP Foundation, and support from Mayo Foundation. Study data included samples collected through the Sun Health Research Institute Brain and Body Donation Program of Sun City, Arizona, USA. The Brain and Body Donation Program has been supported by the National Institute of Neurological Disorders and Stroke (U24 NS072026 National Brain and Tissue Resource for Parkinson’s Disease and Related Disorders), the National Institute on Aging (P30 AG019610 and P30AG072980, Arizona Alzheimer’s Disease Center), the Arizona Department of Health Services (contract 211002, Arizona Alzheimer’s Research Center), the Arizona Biomedical Research Commission (contracts 4001, 0011, 05-901 and 1001 to the Arizona Parkinson’s Disease Consortium) and the Michael J. Fox Foundation for Parkinson’s Research. We would like to thank John Q. Trojanowski (deceased) for his leadership at the Center for Neurodegenerative Disease Research, which helped make acquiring samples from University of Pennsylvania Integrated Neurodegenerative Disease Brain Bank possible. Additional support for these studies was provided by the NINDS grant R01-NS080820 (NET), NIA grant R01-AG061796 (NET), NIA grant U19-AG074879 (NET), and Alzheimer’s Association Zenith Fellows Award (NET). We thank the Mayo Clinic Genome Analysis Core (GAC), Co-Directors Julie M. Cunningham, PhD and Eric Wieben, PhD, and supervisor Julie Lau, for their collaboration in the collection of omics data.

## Consent Statement

This study was approved by the Institutional Review Board at Mayo Clinic. All participants or next-of-kin provided consent.

