## Supplemental Figure 1 for "Bridging the Gap: Multi-Omics Profiling of Brain Tissue in Alzheimer’s Disease and Older Controls in Multi-Ethnic Populations"

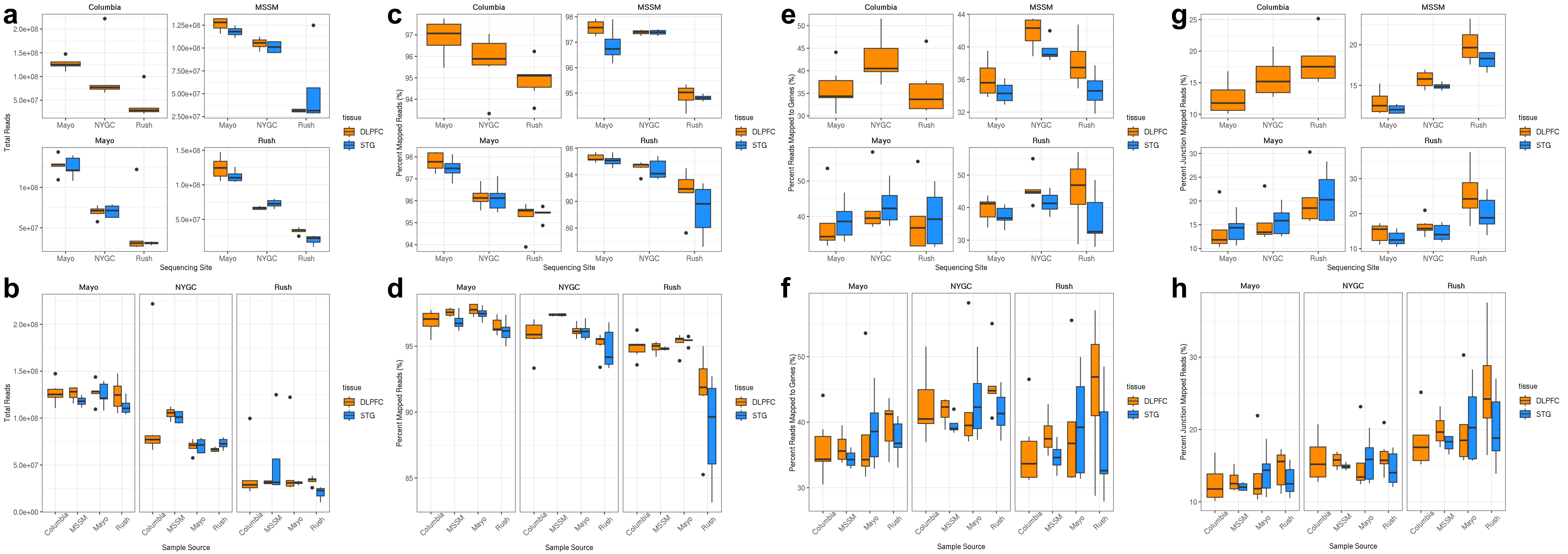


**Supplementary Figure 1. Alignment metrics**. Summaries by tissue sample source (data contribution group) at the top (**a, c, e, g**) and by sequencing site at the bottom (b, d, f and h) are illustrated for the following metrics: Total reads (**a, b**); percent mapped reads (**c, d),** percent reads mapped to genes (**e, f**) and percent junctions (**g, h**).
