## Supplemental Figure 2 for "Bridging the Gap: Multi-Omics Profiling of Brain Tissue in Alzheimer’s Disease and Older Controls in Multi-Ethnic Populations"

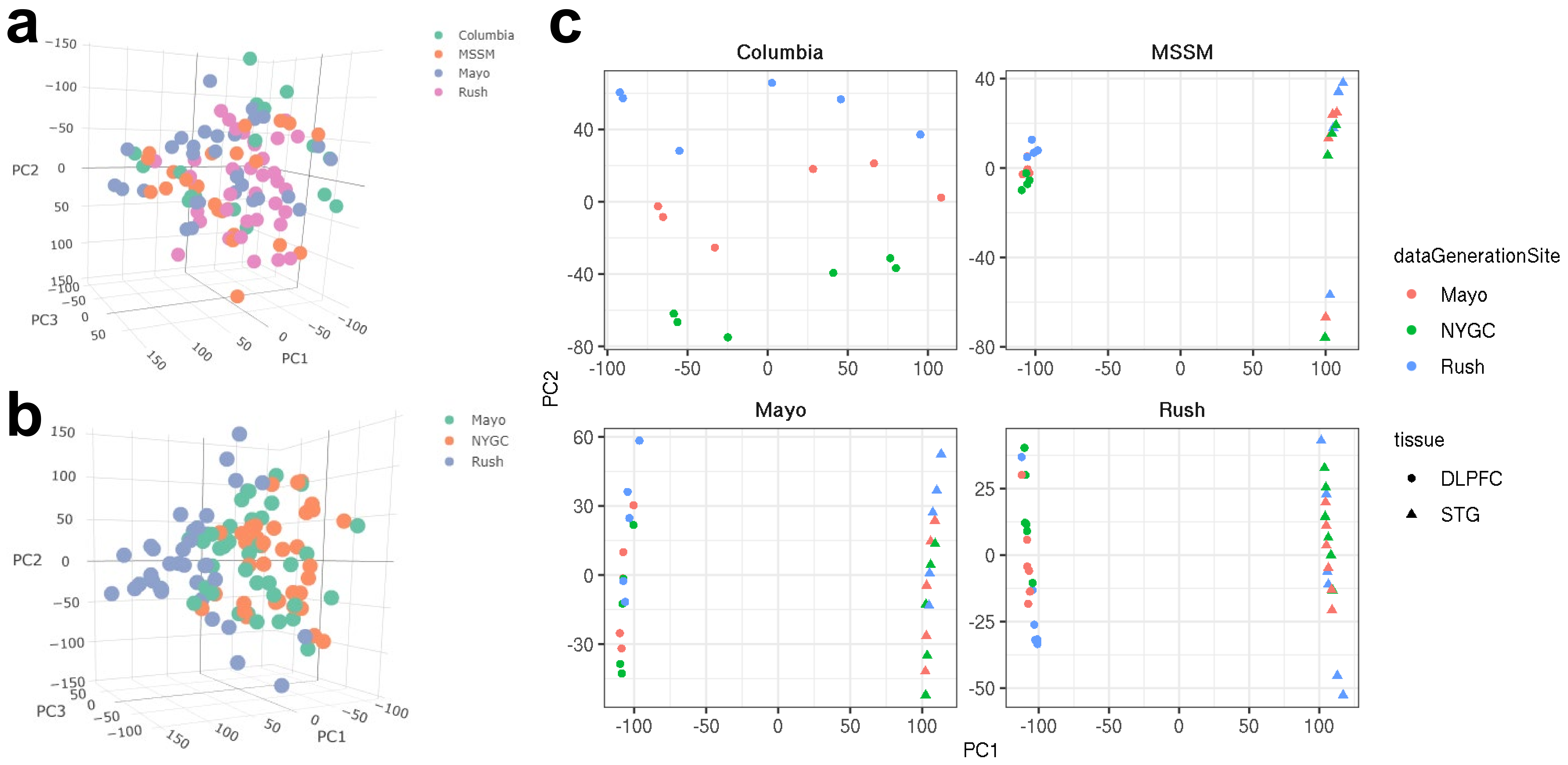


**Supplementary Figure 2. Principal component analysis**. Figure summarizing principal component analysis of gene expression measures. Principal components estimated separately for each tissue and plotted together are shown on the left. The first 3 PCs were plotted and colored either by sample source (**a**) or by sequencing site (**b**). On the right (**c**), PCs were estimated separately for each tissue contribution site and plotted together (first two PCs). On the right, PCs are colored by sequencing site, and the shape of the point represents tissue type.
