## Supplemental Table 1 for "Bridging the Gap: Multi-Omics Profiling of Brain Tissue in Alzheimer’s Disease and Older Controls in Multi-Ethnic Populations"

**Supplementary Table 1. Patient Characteristics of samples sourced from AMP-AD 1.0 to balance proteomics batches**

|  | **AMP-AD 1.0 Individuals** |
| --- | --- |
| **Characteristic** | **N=305** |
| **Cohort** |  |
| *Mt Sinai Brain Bank* | 112 (37%) |
| *ROS* | 145 (48%) |
| *MAP* | 48 (16%) |
| **Female sex, N (%)** | 192 (63%) |
| **Age at death^a^ in years, median (range)** | 87.9  (61-90+) |
| **Race^b^, N (%)** |  |
| *Black or African American* | 3 (1%) |
| *Non-Hispanic White* | 291 (95%) |
| *Other* | 10 (3%) |
| *Missing or unknown* | 1 (0%) |
| **Hispanic ethnicity, N (%)** | 10 (3%) |
| **APOE genotype, N (%)** |  |
| *22* | 1 (0%) |
| *23* | 35 (11%) |
| *24* | 6 (2%) |
| *33* | 147 (48%) |
| *34* | 83 (27%) |
| *44* | 7 (2%) |
| *Missing or unknown* | 26 (9%) |
| **CERAD, N (%)** |  |
| *None/No AD/C0* | 68 (22%) |
| *Sparse/Possible/C1* | 27 (9%) |
| *Moderate/Probable/C2* | 80 (26%) |
| *Frequent/Definite/C3* | 130 (43%) |
| **Braak stage, N (%)** |  |
| *None* | 8 (3%) |
| *I* | 21 (7%) |
| *II* | 23 (8%) |
| *III* | 74 (24%) |
| *IV* | 75 (25%) |
| *V* | 53 (17%) |
| *VI* | 51 (17%) |
| **NIA Reagan** |  |
| No AD | 8 (3%) |
| *Low Likelihood* | 93 (30%) |
| *Intermediate Likelihood* | 118 (39%) |
| *High Likelihood* | 86 (28%) |
| **Derived AD outcome** |  |
| *Control* | 83 (27%) |
| *AD* | 167 (55%) |
| *Other* | 55 (18%) |
